# CRISPR-Mediated Generation and Characterization of a *Gaa* Homozygous c.1935C>A (p.D645E) Pompe Disease Knock-in Mouse Model Recapitulates Human Infantile Onset-Pompe Disease

**DOI:** 10.1101/2022.05.30.494061

**Authors:** Shih-hsin Kan, Jeffrey Y. Huang, Jerry Harb, Allisandra Rha, Nancy D. Dalton, Chloe Christensen, Yunghang Chan, Jeremy Davis-Turak, Jon Neumann, Raymond Y. Wang

**Affiliations:** CHOC Children’s Research Institute, Orange, CA 92868; School of Medicine, New York Medical College, NY 10595; ROSALIND, San Diego, CA, 92126; Transgenic Mouse Facility, University of California Irvine, Irvine, CA 92697; Division of Metabolic Disorders, CHOC Children’s Specialists, Orange, CA 92868; Department of Pediatrics, University of California-Irvine, Irvine, CA 92697

## Abstract

Pompe disease (PD) is an autosomal recessive disorder caused by deficient lysosomal acid α-glucosidase (GAA), leading to reduced degradation and subsequent accumulation of intra-lysosomal glycogen in tissues, especially skeletal and oftentimes cardiac muscle. The c.1935C>A (p.Asp645Glu) variant is the most frequent *GAA* pathogenic mutation in people of Taiwanese and Southern Chinese ethnicity, causing infantile-onset PD (IOPD), which presents neonatally with severe hypertrophic cardiomyopathy, profound muscle hypotonia, and respiratory failure leading to premature death if untreated.

To further investigate the pathogenic mechanism and facilitate development of therapies pertaining to this variant, we applied CRISPR-Cas9 homology-directed repair (HDR) using a novel dual sgRNA approach flanking the target site to generate a *Gaa^Em1935C>A^* knock-in mouse model as well as a myoblast cell line carrying the *Gaa* c.1935C>A mutation. Herein we describe the molecular, biochemical, physiological, histological, and behavioral characterization of 3-month-old homozygous *Gaa^Em1935C>A^* mice.

Homozygous *Gaa^Em1935C>A^* knock-in mice exhibited normal *Gaa* mRNA expression levels relative to wild-type mice, but GAA enzymatic activity was almost completely abolished, leading to a substantial increase in tissue glycogen storage, and significant concomitant impairment of autophagy. Echocardiography of 3-month-old knock-in mice revealed significant cardiac hypertrophy. The mice also demonstrated skeletal muscle weakness but, paradoxically, not early mortality. Longitudinal studies of this model, including assessment of its immune response to exogenously supplied GAA enzyme, are currently underway.

In summary, the *Gaa^Em1935C>A^* knock-in mouse model recapitulates the molecular, biochemical, histopathologic, and phenotypic aspects of human IOPD caused by the *GAA* c.1935C>A pathogenic variant. It is an ideal model to assess innovative therapies to treat IOPD, including personalized therapeutic strategies that correct pathogenic variants, restore GAA activity and produce functional phenotypes.

## Introduction

Glycogen storage disease type II, also called Pompe disease (PD; OMIM#232300), is an autosomal recessive disorder resulting from malfunction of lysosomal acid α-glucosidase (GAA; EC 3.2.10.20) caused by mutations in the *GAA* gene (OMIM#606800). GAA deficiency leads to reduced glycogen degradation and accumulation of intra-lysosomal glycogen with pronounced glycogen storage in cardiac and skeletal muscle. Increased glycogen storage in myocytes, brain, and spinal cord anterior horn neurons results in muscle weakness, which varies in age of onset and severity according to the level of residual GAA enzymatic activity[1]. PD presents as a spectrum of phenotypes, typically classified into infantile-onset form (IOPD) and late-onset form (LOPD) based on the time of disease onset[2–4]. Patients with severe IOPD have neonatal onset and a rapidly progressive disease with prominent cardiomyopathy, general muscle weakness and hypotonia, respiratory problems and drastically reduced life expectancy. Patients with LOPD have a more slowly progressive proximal skeletal myopathy eventually resulting in mobility problems and respiratory difficulties, but generally do not present with hypertrophic cardiomyopathy[3].

Recombinant GAA (rhGAA) enzyme replacement therapy (ERT) was developed to treat PD and approved by the FDA in 2006. ERT improves the survival of patients and is very effective at reducing glycogen levels in heart muscle and reversing cardiac symptoms. However, only partial recovery of muscle strength can be achieved with ERT. Surviving children still have glycogen buildup in other muscles and struggle with basic activities such as talking, walking, eating or even breathing[5, 6].

The *GAA* gene has a very heterogeneous mutational spectrum, with more than 900 *GAA* variants documented in the Pompe disease GAA variant database[7–9]. Among these variants, the *GAA* c.1935C>A transversion in exon 14, which results in the p.Asp645Glu (p.D645E) missense mutation, is the most frequent pathogenic variant associated with IOPD in the Taiwanese, Southern Han Chinese, and Southeast Asian populations, but is not frequently reported in any other region[8, 10]. In Taiwanese populations, this c.1935C>A variant represents 36-80% of mutations[11, 12], indicating existence of a founder effect that stems from a diaspora of Southern Han Chinese to Taiwan and other locations[12].

Here, we report the generation of a *Gaa^Em1935C>A^* (p.D645E) knock-in (KI) mouse model of PD by CRISPR-Cas9 homology-directed repair (HDR) using a dual sgRNA approach. The primary objective of this study is to characterize the molecular, biochemical, physiological, histological, and behavioral phenotypes of this KI mouse model. We anticipate that this novel *Gaa^Em1935C>A^* mouse model will be a valuable research tool, especially when compared to other *Gaa* knockout and KI models. Altogether, preclinical KI models of PD will further accelerate our understanding of how pathogenic *GAA* mutations result in variable disease onset, progression, and response to current and future therapeutic strategies.

## Results

### *Gaa^c.1935^* target locus guide RNA and donor ssODN design

*In silico* design of CRISPR-Cas9 guide RNAs (gRNAs) specific for the *Gaa^c.1935^* target locus was performed using Genetic Perturbation Platform (GPP) sgRNA Designer[13]. Candidate gRNAs were selected using the following criteria: 1) top combined rank score (based on on-target efficacy and off-target specificity scores) and 2) proximity of predicted Cas9 nuclease cut site to the *Gaa^c.1935^* target locus. Further potential gRNA off-target analysis was performed using Genome Target Scan (GT-Scan)[14]. Three gRNAs were used in this study to generate *Gaa^c.1935C>A^* KI C2C12 cells and transgenic mice: gRNA-1 (5’-CGCAGATGTCCGCCCCGACC-3’), gRNA-2 (5’-GCAGATGTCCGCCCCGACCA-3’), and gRNA-3 (5’-GGGCGTGCCCCTGGTCGGGG-3’).

*Gaa^c.1935^* gRNA-1 and gRNA-2 expression vectors and their respective single-stranded donor oligonucleotides (ssODN) were electroporated into C2C12 mouse myoblasts to assess *in vitro* on-target editing and HDR efficiency. *Gaa^c.1935^* gRNA-2 demonstrated higher on-target editing (26.7±10.7%) and HDR efficiency (5.4±3.4%) than gRNA-1 (on-target editing: 13.2±3.7%; HDR efficiency: 3.8±0.6%) (Table 1). Given our prior success in generating *Gaa^Em1826dupA^* KI cell and transgenic mouse lines using a dual overlapping gRNA strategy[15], we applied this same method towards the generation of *Gaa^c.1935C>A^* KI C2C12 cells and *Gaa^Em1935C>A^* mice.

**Table 1:**
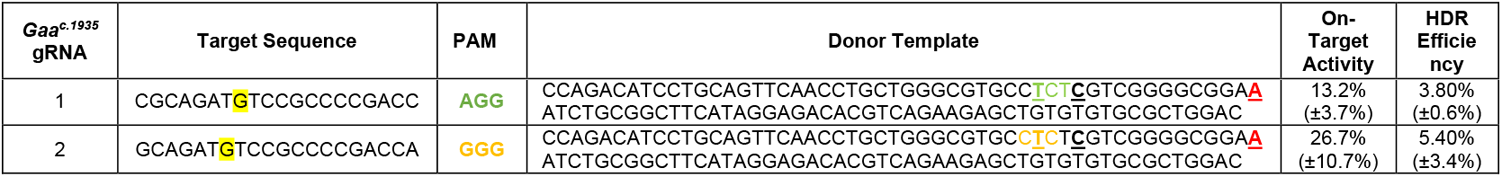
*Gaa^c.1935^* guide RNA on-target activity and HDR efficiency. The target sequence, PAM motifs and donor templates used for testing of *Gaa^c.1935^* guide RNAs in C2C12 mouse mybloasts are outlined. *Gaa^c.1935^* locus for each gRNA target sequence is highlighted in yellow. Desired *Gaa^c.1935^* KI mutation (red), silent PAM site (either green or gold, corresponding to gRNA) and gRNA seed region (black) mutations are bolded and underlined in the donor template sequence. Total on-target Cas9 nuclease activity and HDR efficiency for each *Gaa^c.1935^* guide RNA condition is displayed as the average of two independent experiments.

### Generation and characterization of *Gaa^c.1935C>A^* KI C2C12 cell line

Using the dual overlapping gRNA strategy, we were able to successfully isolate and expand a *Gaa^c.1935C>A^* KI C2C12 clonal line following puromycin-resistant selection of cells electroporated with *Gaa^c.1935C^* gRNA-1 and gRNA-2 and their respective donor ssODN (Table 1; Fig. 1A). Sanger sequence results confirmed that the *Gaa^c.1935C>A^* KI mutation along with a *Gaa^c.1920C>T^* silent PAM mutation were successfully introduced into the clonal line (Fig. 1B).

**Figure 1.**
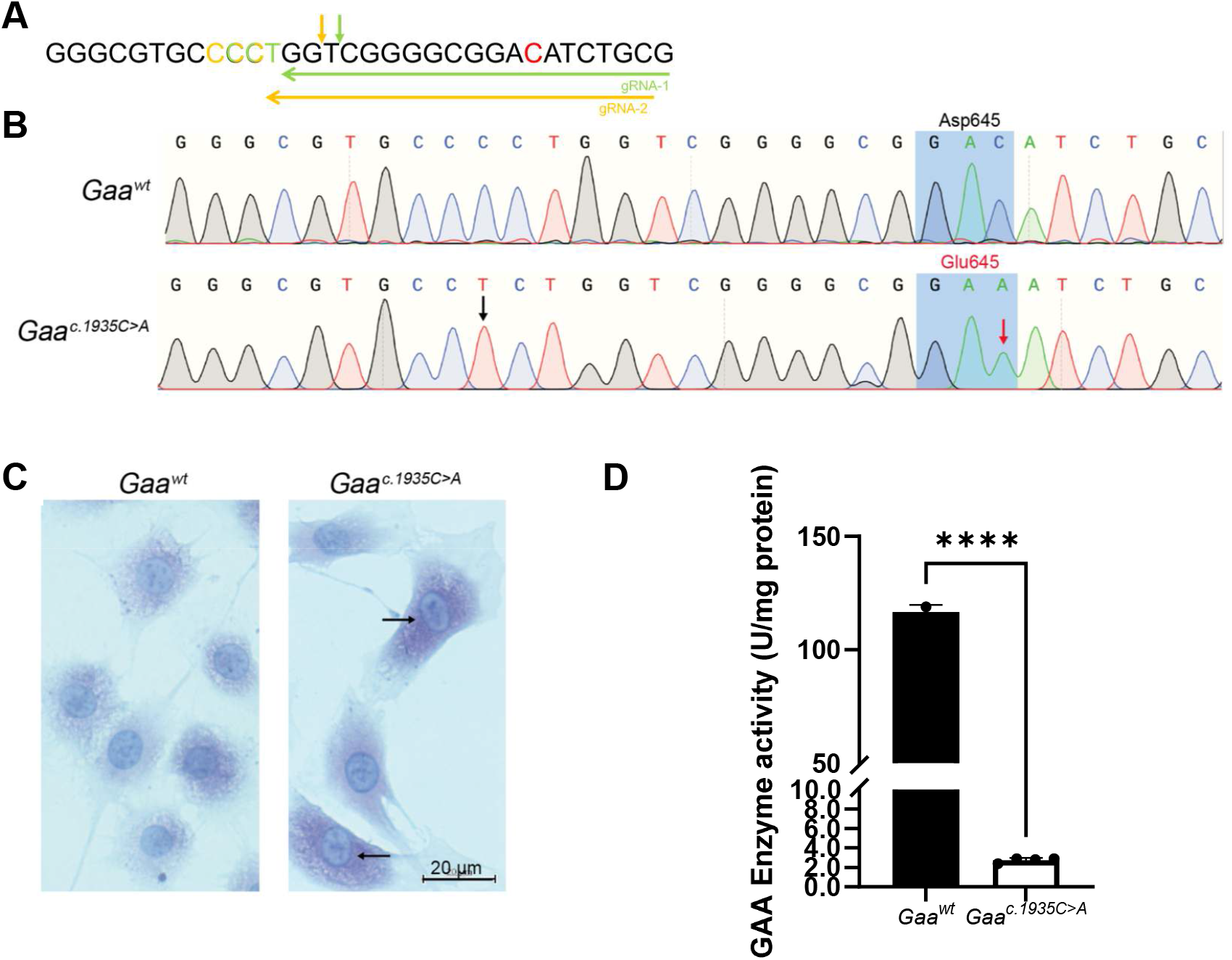
Generation and characterization of a *Gaa^c.1935C>A^* C2C12 myoblast clonal cell line. (A) Sequences of guide RNAs targeting the *Gaa^c.1935^* target locus. Horizontal arrow indicates antisense guide RNAs used in this study. Protospacer adjacent motifs (PAM; NGG) are highlighted in color corresponding to the respective guide RNA. The *Gaa^c.1935^* locus for targeted cytosine to adenine transversion is highlighted in red. (B) Sanger sequencing chromatograms of controls (*Gaa^wt^*) and clonal KI (*Gaa^c.1935C>A^*) C2C12 myoblast genomic DNA at the *Gaa^c.1935^* locus. Black arrow indicates a silent mutation at the PAM site (*Gaa^c.1920C>T^*). Red arrow indicates the desired KI mutation (*Gaa^c.1935C>A^*). Gray shaded region indicates amino acid change from aspartic acid (Asp; GAC) to glutamic acid (Glu; GAA) at position 645. (C) Periodic-acid Schiff (PAS) staining of control (*Gaa^wt^*) and clonal KI (*Gaa^c.1935C>A^*) C2C12 myoblasts. Fixed cells were stained by PAS staining (purple-magenta) and counterstained by hematoxylin (blue). Only *Gaa^c.1935C>A^* KI myoblasts display accumulated PAS staining (see arrows). Representative images were captured on a bright-field microscope (Olympus) at 40x objective magnification. (D) GAA enzymatic activity in *Gaa^wt^* and *Gaa^c.1935C>A^* C2C12 myoblasts. Very low GAA activity (~2.3%) was measured in *Gaa^c.1935C>A^* C2C12 myoblasts compared to *Gaa^wt^* C2C12 myoblasts. GAA enzymatic activity was measured using a fluorometric 4-MU α-D-glycoside assay and normalized to total amount of sample protein. Data generated from three independent experiments are shown as mean ± SD. Comparisons were analyzed with unpaired one-tailed *t*-tests. \*\*\*\**p*<0.0001.

In comparison to *Gaa^wt^* cells, *Gaa^c.1935C>A^* KI cells displayed increased PAS staining, indicating the accumulation of glycogen (Fig. 1C). Furthermore, GAA enzymatic activity was almost abolished in *Gaa^c.1935C>A^* KI cells relative to *Gaa^wt^* cells; less than 2.3% of WT GAA activity was detected in the KI cell line (Fig. 1D). Taken together, these results demonstrate that our *Gaa^c.1935C>A^* KI C2C12 cell line exhibits a molecular and biochemical phenotype observed in human PD and can be utilized as an *in vitro* model for further study.

### Generation and characterization of *Gaa^Em1935C>A^* transgenic mice

We next applied the dual overlapping gRNA strategy *in vivo* using *Gaa^c.1935^* gRNA-2 and gRNA-3 and their respective donor ssODN (Fig. 2A, B). We were able to successfully generate *Gaa^Em1935C>A^* KI mice via pronuclear injection of C57BL/6NJ single-cell embryos by standard methods[16]. Use of dual overlapping gRNA achieved a high percentage (89.7%) of on-target editing activity in genome-edited founder mice with any *Gaa* mutation as well as 38.5% HDR efficiency (founder mice positive for the desired *Gaa^Em1935C>A^* KI mutation) (Table 2). For mating and segregation of the *Gaa^Em1935C>A^* KI mutation, we genotyped founder mice and selected those with the lowest levels of mosaicism for the wild-type allele as determined by Sanger sequencing and WGS analyses. Sanger sequencing analysis confirmed presence of the *Gaa^c.1935C>A^* KI mutation along with silent PAM and seed region mutations in *Gaa^Em1935C>A^* founder mice (Fig. 2C). To further determine the extent of genomic mosaicism in our *Gaa^Em1935C>A^* founder mice, we performed WGS at >50x coverage and on-target locus alignment analysis of *Gaa^c.1935C>A^* founder and *Gaa^wt^* mice. For on-target analysis, *Gaa^c.1935^* target loci from aligned FASTQ reads were designated to four categories: *Gaa^c.1935^* KI mutation; insertion/deletion mutation (indel); no mutation; and nonspecific mutation. WGS analysis demonstrated highly efficient integration of the desired *Gaa^c.1935C>A^* KI mutation (>50% for *Gaa^c.1935C>A^* founder #1 and >25% for *Gaa^c.1935C>A^* founder #2) with indel and nonspecific mutations comprising a minority of genomic editing events in *Gaa^c.1935C>A^* founder mice selected for mating (Fig. 2D). For off-target analysis of the WGS data, the only result was the intended *Gaa^c.1935C>A^* mutation. Five genomic regions were predicted by GT-Scan as potential off-target sites of the CRISPR sgRNAs, but no SNVs were detected within 500 bp of these sites.

**Figure 2.**
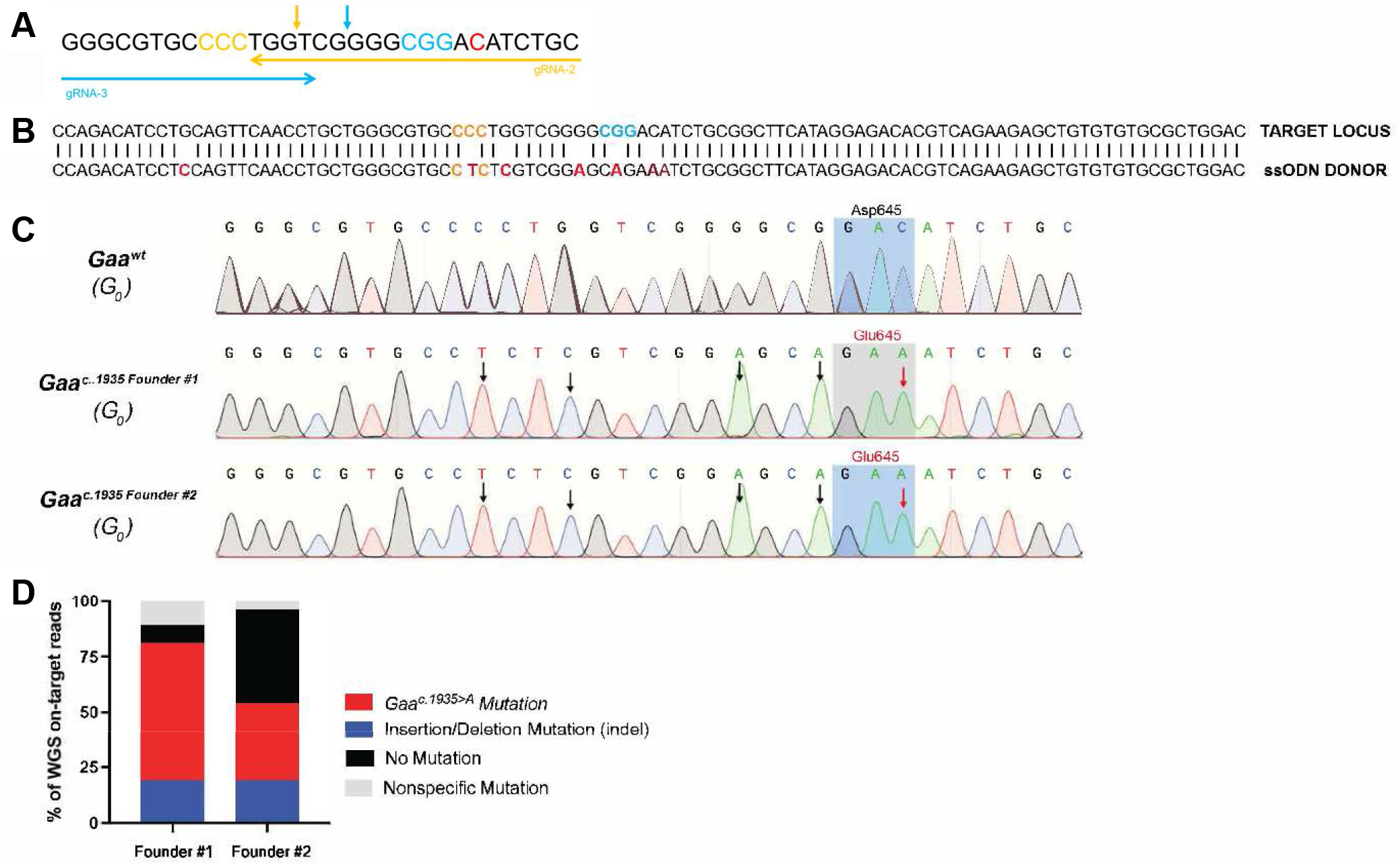
Generation of a *Gaa^Em1935C>A^* transgenic mouse line. (A) Dual overlapping guide RNA approach targeting the *Gaa^c.1935^* target locus. Arrowhead direction indicates whether guide RNA is sense (right) or antisense (left). PAM sequences (NGG) are highlighted in color corresponding to the guide RNA arrow. The *Gaa^c.1935^* locus for target adenine to cytosine nucelotide transversion is highlighted in red. Expected Cas9 nuclease cut sites are shown as vertical arrows in color corresponding to the guide RNA arrow. (B) Sequence of single-stranded donor oligonucleotide (ssODN) for targeted integration of the *Gaa^Em1935C>A^* KI mutation. PAM motifs are indicated in gold (gRNA-2) or blue (gRNA-3). Intended synonymous variants at PAM sites (*Gaa^c.1920C>T^, Gaa^c.1932G>A^*), gRNA seed regions (*Gaa^c.1923G>C^, Gaa^c.1929G>A^*), a restriction fragment length polymorphism (RFLP) site (*Gaa^c.1896G>C^*), and the desired KI mutation are highlighted in red. (C) Sequencing chromatograms of control (*Gaa^wt^*), founder #1 (*Gaa^c.1935 Founder #1^*), and founder #2 (*Gaa^c.1935 Founder #2^*). Black arrows indicate synonymous variant edits at PAM sites (*Gaa^c.1920^, Gaa^c.1932^*) or gRNA seed regions (*Gaa^c.1923^, Gaa^c.1929^*). Red arrows indicate the desired KI mutation (*Gaa^c.1935C>A^*). Gray shaded region indicates amino acids at position 645 for each mouse. (D) WGS on-site target analysis (>50x read depth) of the *Gaa^c.1935^* locus in G_0_ founder #1 and G_0_ founder #2. WGS analysis demonstrates highly efficient on-target genome-editing in these founder mice. Data are presented as stacked bar graphs indicating the percentage of WGS on-target reads for each event category.

**Table 2:** Dual overlapping gRNA strategy and outcomes of *Gaa^Em1935C>A^* mouse generation. Dual overlapping gRNA components and concentrations used in pronuclear microinjection of C57BL/6NJ fertilized zygotes are outlined. Each crRNA was hybridized with tracrRNA at a 1:1 ratio to form gRNA duplexes. Equimolar amounts of gRNAs were then combined with 3xNLS spCas9 at a 1:1 ratio to form the RNP complex. Numbers of founder mice positive for any *Gaa* mutation and founder mice with the *Gaa^c.1935^* KI mutation are reported as percentages.

| Dual gRNA Strategy Components | Concentration | Founder Mice with any *Gaa* mutation (% Positive) | Founder Mice with *Gaa*<sup>Em1935C>A</sup> mutation (% Positive) |
| --- | --- | --- | --- |
| *Gaa*<sup>c.1935</sup> crRNAs (gRNA-2, gRNA-3) | 3 µM<br>RNP complex | 89.7%<br>(35 of 39 founders) | 38.5%<br>(15 of 39 founders) |
| tracrRNA |  |  |  |
| 3xNLS spCas9 |  |  |  |
| Donor ssODN | 10 ng/µL |  |  |

### *Gaa^Em1935C>A^* KI mice have severe GAA enzymatic deficiency and glycogen storage in cardiac, skeletal muscle, and brain tissue

The missense *Gaa^c.1935C>A^* mutation in exon 14 of the *Gaa* gene leads to an amino acid substitution; therefore, nonsense mediated decay is not expected in mRNA transcripts carrying the *Gaa^c.1935C>A^* mutation. The ΔC_t_ between mouse *Gaa* and housekeeping gene *Gapdh* acquired by RT-PCR among WT, HET, and KI groups are almost identical, suggesting this *Gaa^c.1935C>A^* mutation does not affect *Gaa* gene expression (Fig. 3A).

**Figure 3.**
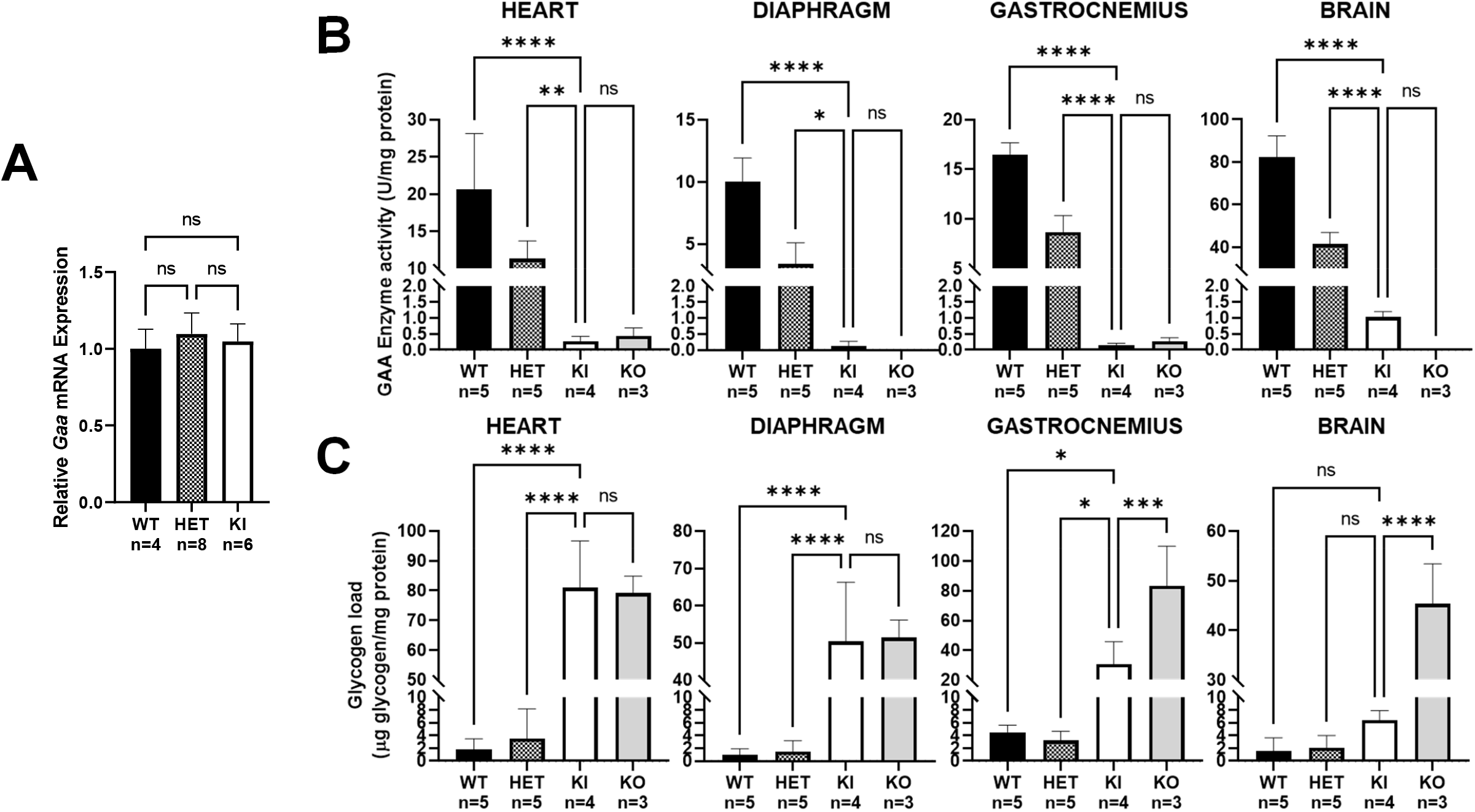
Molecular and biochemical characterization of *Gaa^c.1935C>A^* KI mice. (A) *Gaa* mRNA expression in tail or liver biopsy samples from 3-month-old WT (n=4; black bar), HET (n=8; dotted bar), and KI (*Gaa^Em1935C>A^*; n=6; white bar) mice. No significant difference in *Gaa* mRNA transcript expression was detected among WT, HET, and KI samples. *Gaa* expression levels were measured by TaqMan probe-based quantitative real-time PCR using the ΔC_t_ method for comparison of the target gene (*Gaa*) to the reference gene (*Gapdh*). (B) GAA enzyme activity in heart, diaphragm, and gastrocnemius muscle tissues and brain homogenate from WT (n=5; black bars), HET (n=5; dotted bars), KI (*Gaa^Em1935C>A^*; n=4; white bars), and KO (*Gaa^tm1Rabn^*; n=3; grey bars) mice was measured using a fluorometric 4-MU α-D-glucopyranoside assay and normalized to the amount of sample protein. (C) Glycogen level was measured in the same tissues used for analysis in (B) using a colorimetric assay. KO mice displayed significantly elevated glycogen levels relative to WT and HET mice in all tissues assayed. However, KI mice showed a significant elevation of glycogen levels in muscle tissues, but no significant elevation in brain. The amount of glycogen was normalized to the amount of sample protein. Data were generated from at least three independent experiments and shown as mean ± SD. All comparisons were analyzed using one-way ANOVA with the Tukey post-hoc test. \**p*<0.05, \*\**p*<0.01, \*\*\**p*<0.001, \*\*\*\**p*<0.0001. ns: not significant.

GAA enzymatic activity was measured with artificial fluorometric 4-MU substrate as described previously[15]. The results were consistent with the other findings from this study, showing that the HET group had close to 50% of the level of enzymatic activity observed in the WT group in each muscle tissue and brain tissue sample tested, indicating that the one WT allele produced functional enzyme, but not the c.1935C>A allele. Compared to tissue from WT or HET animals, tissue from KI (*Gaa^Em1935C>A^*) and KO (*Gaa^tm1Rabn^*/J; exon 6 knock-out) animals had significantly decreased in GAA enzymatic activity (about 1% of WT levels) (Fig. 3B).

Compared to the unaffected WT or HET groups, KI and KO mice had abnormally elevated lysosomal glycogen storage in heart, diaphragm, and gastrocnemius muscle tissue. Interestingly, the increase in glycogen storage in whole-brain homogenate was observed in KO mice, but not in KI mice, which had a slight, but not statistically significant, increase in glycogen load (Fig. 3C).

### *Gaa^Em1935C>A^* KI mice have impaired muscle autophagy

Excessive autophagic buildup is well-documented in PD patients and in PD mice[17, 18] and may be a potential mechanism of pathogenesis. Microtubule-associated protein light chain 3 (LC3B) is a protein component of autophagosomes, which are quickly degraded under normal physiological conditions and are hardly detectable. Cleavage of LC3B at the carboxy terminus immediately following synthesis yields the cytosolic LC3B-I form. LC3B-I is converted to LC3B-II when autophagic processes are activated; the LC3B-II:LC3B-I ratio is therefore used as an indicator of autophagy. To examine the autophagy status of the *Gaa^Em1935C>A^* KI mouse, western blotting for LC3B was performed using tissue homogenate (Fig. 4A). Compared to WT and HET mice, both KI and KO mice showed significantly increased levels of both LC3B-I and LC3B-II in gastrocnemius, with striking increases in LC3B-II (Fig. 4B, 4C). When comparing the ratio of LC3B-II to total LC3B, KI mice showed a significant increase in the three muscle tissues examined, but no increase was observed in brain homogenate (Fig. 4D).

**Figure 4.**
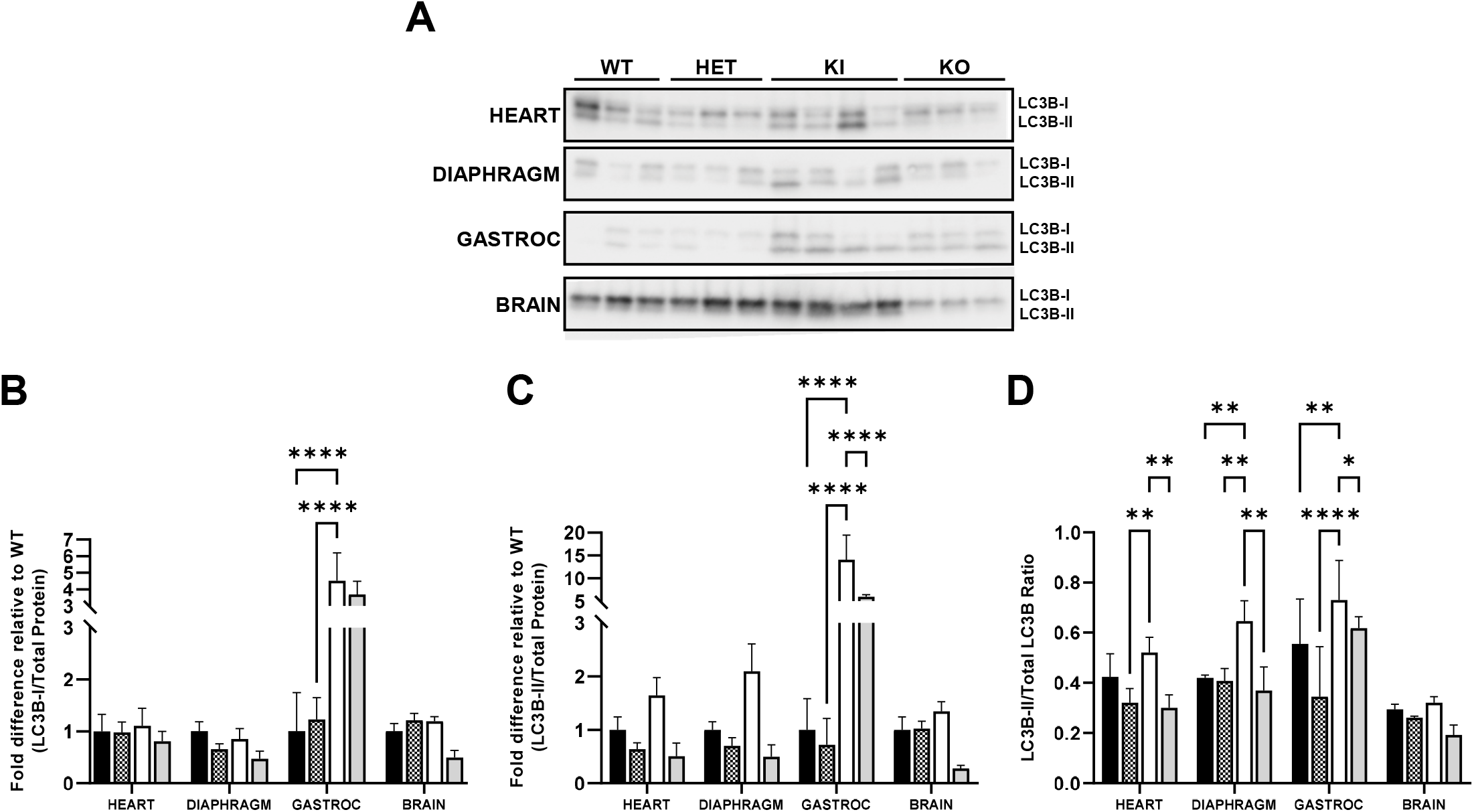
Autophagy impairment in the *Gaa^c.1935C>A^* KI mouse model. (A) Representative western blot images of autophagy-associated proteins (LC3B-I and LC3B-II) from tissue homogenate from heart, diaphragm, gastrocnemius (GASTROC), and brain of WT (n=3; black bars), HET (n=3; dotted bars), KI (n=4; white bars), and *Gaa^tm1Rabn^* (KO, n=3; grey bars) mice. Prominent LC3B-II bands can be seen in KI and KO tissues. (B) LC3B-I and (C) LC3B-II protein levels normalized to the amount of total protein. (D) LC3B-II normalized to the total LC3B ratio. LC3B protein intensity was quantified by densitometric analysis of the western blots, and the amount of total protein was determined by densitometric analysis of stain-free gels. Data were generated from at least three independent western blots, and values are shown as mean ± SD. All comparisons were analyzed using one-way ANOVA with the Tukey post-hoc test. \**p*<0.05, \*\**p*<0.01, \*\*\*\**p*<0.0001.

### *Gaa^Em1935C>A^* KI mice display left ventricular cardiac hypertrophy at 3 months of age

Neonatal-onset hypertrophic cardiomyopathy is a common clinical presentation in patients with IOPD. To explore the anatomical features and physiological function of hearts in the KI mice, echocardiography was performed on 3-month-old mice. M-mode images obtained by echocardiography were used to measure multiple parameters including wall thickness, internal diameter, and heart rate. Many additional functional parameters can be derived from these measurements to determine temporal left ventricular (LV) wall motion as an index for LV contractile patterns and chamber size (Fig. 5A).

**Figure 5.**
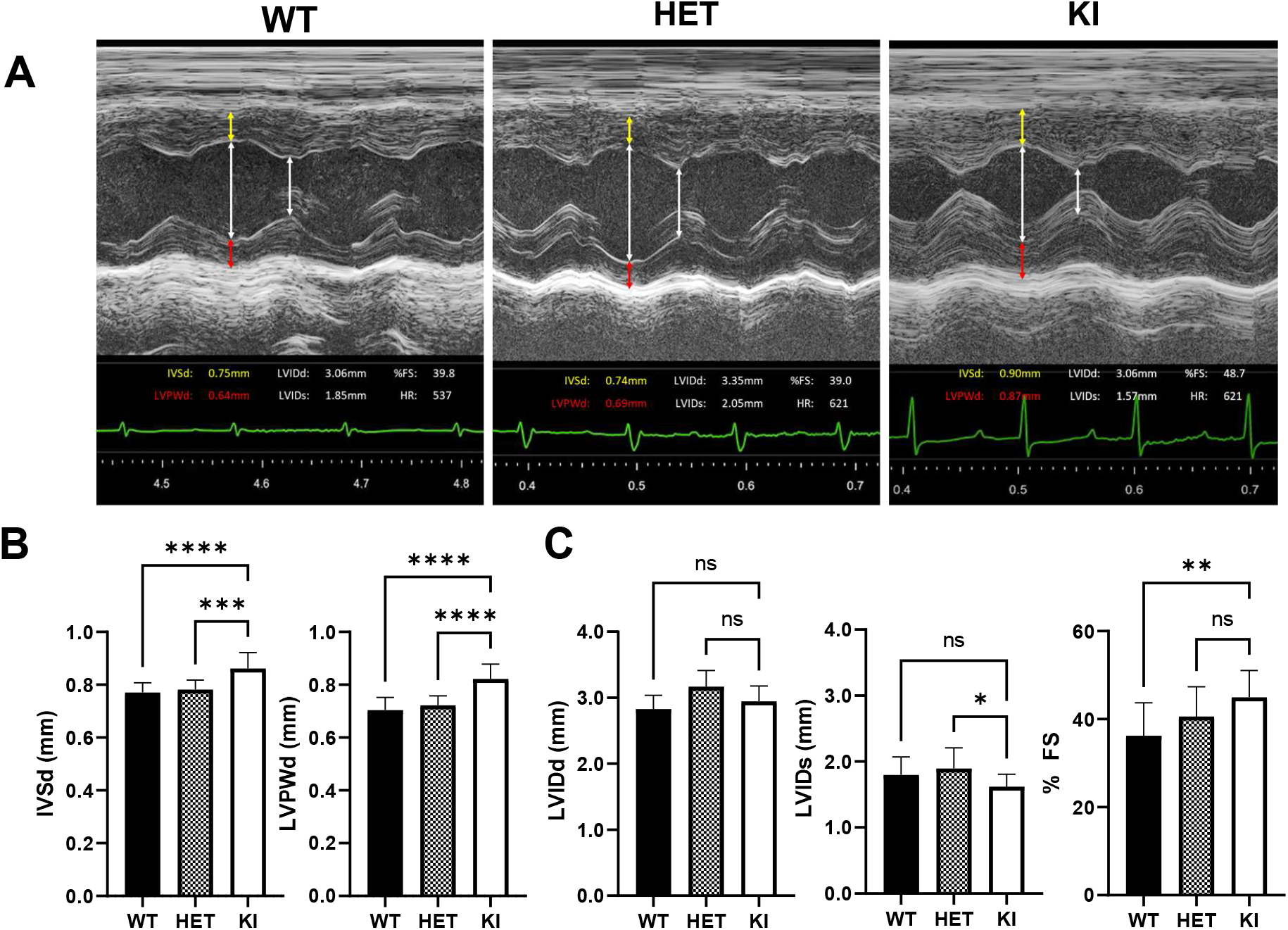
Three-month-old *Gaa^Em1935C>A^* KI mice display anatomical features of left ventricular cardiac hypertrophy. (A) Representative M-mode echocardiographic images showing cardiac dimensions: IVSd: yellow arrows; LVPWd: red arrows, and LVIDd / LVIDs: white arrows. (B) Comparison of ventricular dimension measurements: IVSd and LVPWd. (C) Comparison of myocardial contraction measurements: LVIDd and LVIDs. (D) Comparison of calculated fractional shortening and LVMI. All measurements from WT (n=12; black bars), HET (n=10; dotted bars), and KI (*Gaa^.Em1935C>A^*; n=10; white bars) mice were obtained from 3-month-old mice. Data are shown as mean ± SD. Heart rate (HR) was maintained greater than 500 bpm throughout measurements. All comparisons were analyzed using one-way ANOVA with the Tukey post-hoc test. \**p*<0.05, \*\**p*<0.01, \*\*\**p*<0.001, \*\*\*\**p*<0.0001. ns: not significant.

Increases in interventricular septal diameter (IVSd), LV posterior wall diameter (LVPWd), and LV mass index (LVMI) were observed in KI mice, compared to WT and HET mice (Fig. 5B), indicating pronounced hypertrophic cardiomyopathy. Measurements of myocardial contraction showed a slight decrease in LV systolic internal diameter (LVIDs) in the KI mouse, but no significant difference in LV diastolic internal diameter (LVIDd) was observed among WT, HET, and KI mice (Fig. 5C). Increased fractional shortening indicative of cardiac contractile dysfunction was observed in KI mice (Fig. 5D). Echocardiographic data therefore indicates early hypertrophic cardiomyopathy phenotypes in 3-month-old *Gaa^c.1935C>A^* KI mice. The data presented in Fig. 5 show no gender differences in these parameters (Supplementary Fig. 1).

### Reduced forelimb grip strength in *Gaa^Em1935C>A^* KI mice

The forelimb grip strength test is commonly used to evaluate neuromuscular dysfunction in mice by measuring the deterioration of skeletal muscle. Peak tension force was recorded as the mice lost their grip on the force transducer bar and normalized to bodyweight for analysis by gender group.

First, mouse body weight is known to differ between genders at 3 months of age[19]. The mean ± SD body weights of male and female mice in our study cohort were 28.73 ± 3.13 g and 21.76 ± 2.36 g, respectively. In each gender cohort, there was no significant difference in bodyweight across WT, HET, and KI mice (Fig. 6). In addition, at 3 months of age, the male *Gaa^Em1935C>A^* KI mouse showed a significant reduction (~19%) in normalized peak tension force compared to WT mice, indicating decreased forelimb muscle strength in KI mice (Fig. 6). This reduction was observed only in male KI mice, but not in female KI mice.

**Figure 6.**
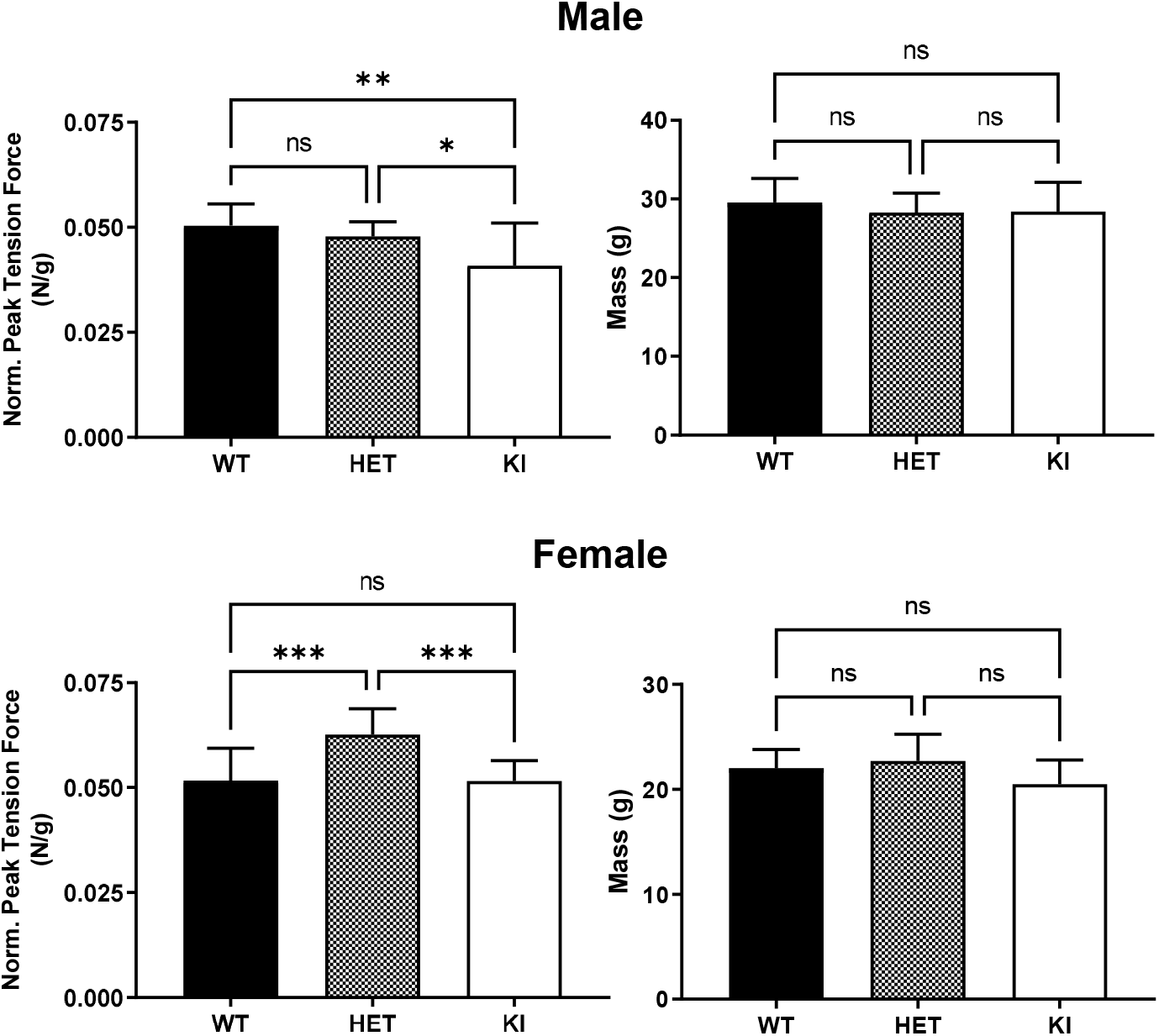
Reduced forelimb grip strength in male *Gaa^Em1935C>A^* transgenic mice. Forelimb peak tension force and body mass measurements in 3-month-old male WT (n=12; black bar), HET (n=12; dotted bar), and KI (n=14; white bar) mice (top panel) and female WT (n=12; black bar), HET (n=12; dotted bar), and KI (n=11; white bar) mice (bottom panel). Forelimb peak tension force was measured using a grip strength meter and taken as the average of 9 trials over 3 days. Data are shown as mean ± SD. All comparisons were analyzed using one-way ANOVA with the Tukey post-hoc test. \**p*<0.05, \*\**p*<0.01, \*\*\**p*<0.001. ns: not significant.

### *Gaa^Em1935C>A^* KI mice show increased muscle glycogen content

PAS staining is routinely used to demonstrate abnormal carbohydrate accumulation in muscle tissue[20]. PAS staining was performed in different muscle tissues (heart, diaphragm, and gastrocnemius) as well as brain from 3-month-old KI mice. Scattered red to magenta PAS staining particles representing the accumulation of glycogen were observed in all three muscle tissue types in the KI mice, but not in WT animals (Fig. 7A). PAS staining with diastase (PAS-D), an enzyme that digests only glycogen, was also applied to consecutive slides to confirm that the particles consisted of glycogen. A decrease in red/magenta signal confirms that excessive accumulation products in tissues comprised only glycogen (Supplementary Fig. 2). The intensity of PAS staining was quantified with the color deconvolution function in ImageJ software[21], and the OD was calculated as described above. Quantified PAS staining showed significant glycogen storage in KI heart and diaphragm, as well as increased gastrocnemius glycogen in KI mice (p=0.06) (Fig. 7A, bottom).

**Figure 7.**
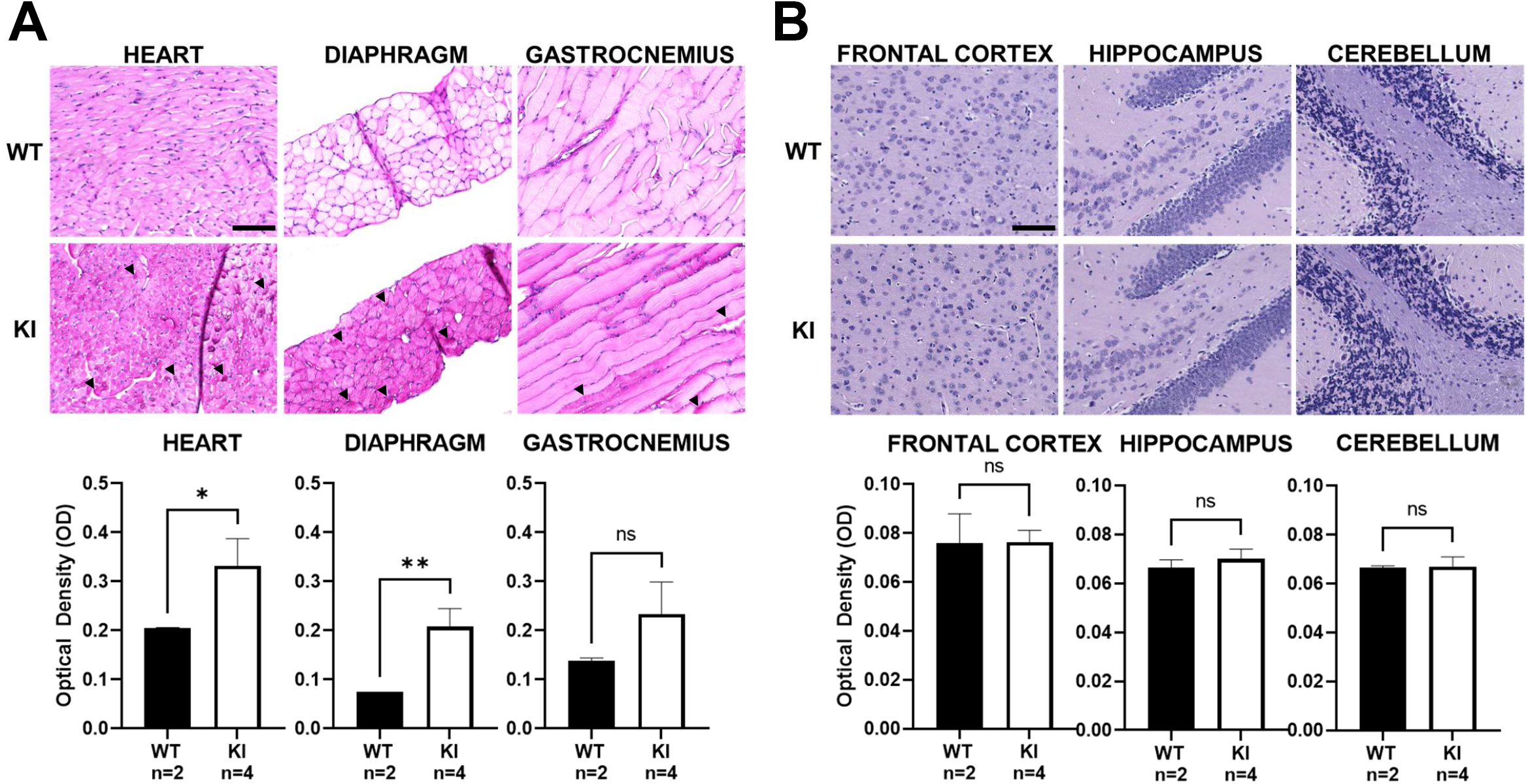
Tissue pathology and glycogen storage in *Gaa^Em1935C>A^* KI mice. Representative images of heart, diaphragm, and gastrocnemius sections from 3-month-old WT and KI mice, stained with hematoxylin/PAS. (A) Areas of abnormal glycogen accumulation (arrowheads) in cardiac and skeletal muscle tissues were observed in KI mice compared to WT mice (top). Quantification of the OD of tissue PAS staining by color deconvolution analysis is shown (bottom). Increased PAS staining signal in KI mice compared to WT mice was detected in all three types of muscle tissue examined. (B) No significant difference in PAS staining between WT and KI mice was observed in brain sections. Three representative brain areas (frontal cortex, hippocampus and cerebellum) are shown (top), with no observed difference between WT and KI mice. Scale bar represents 100 μm. Measurements were obtained from samples from 3-month-old WT (n=12), HET (n=10), and KI (n=10) mice. Data are shown as mean ± SD. All comparisons were analyzed using the unpaired two-tailed *t*-test. \**p*<0.05, \*\**p*<0.01. ns: not significant.

For glycogen storage in brain, three representative areas of brain (frontal cortex, hippocampus, and cerebellum) were examined (Fig. 7B, top). No strong PAS staining was observed in either WT or KI mice, in any of the three brain areas. Quantified PAS staining also demonstrated low PAS OD in brain, with no significant difference between WT and KI brain sections (Fig. 7B, bottom).

In summary, histopathology results showed that the KI mice display early pathological glycogen accumulation in muscle tissues, which is analogous to muscle pathology in IOPD patients.

## Discussion

In southern China and Taiwanese populations, the *GAA* c. 1935C>A (p.Asp645Glu) mutation represents 36%-80% of mutations [11, 12, 22] in IOPD patients. We have successfully applied CRISPR/Cas9 genome editing to install the *Gaa c. 1935C>A* mutation in a mouse myoblast C2C12 cell line and create a novel *Gaa^Em.1935C>A^* KI mouse model; each of which represents a valuable resource for studying IOPD. The KI C2C12 line demonstrates severe GAA enzyme deficiency and glycogen accumulation; the KI mouse model successfully recapitulates molecular, biochemical, histologic, and phenotypic aspects of human IOPD.

While no phenotypic differences were noted between c.1935C>A HET and WT aside from the expected 50% reduction in HET GAA enzymatic activity, the homozygous KI mice demonstrated a significant, PD-like phenotype. KI mice had normal *Gaa* mRNA levels with significantly reduced, but notable level of GAA hydrolysis activity (about 1% of WT) in heart and skeletal muscle, as well as brain tissue. This aligns with observed levels of low GAA enzyme activity (0.08-0.82% of normal range for control) previously measured in homozygous c.1935C>A patient fibroblasts[23]. Significant increases in glycogen storage were observed in KI mouse muscle tissues, consistent with the human *GAA* c.1935C>A IOPD phenotype. It is interesting to note that glycogen storage in brain tissue from *Gaa^Em.1935C>A^* mice was not affected. Autophagic impairment was noted in cardiac and skeletal muscle tissues, consistent with what is observed in human PD and other murine PD models[24]. *Gaa^Em1935C>A^* mice developed hypertrophic cardiomyopathy at approximately two months of age, which becomes quite marked at three months of age. This muscle weakness phenotype may be due to a combination of sequelae from cardiomyopathy and autophagic impairment. Studies are ongoing to assess the life span, natural history, and phenotypic progression of the model.

A significant divergence of the model from human c.1935C>A IOPD is the lack of infantile mortality in KI mice. This KI mouse, along with the *Gaa^Em1826dupA^* KI mouse strain previously generated in our laboratory[15] and other previously published *Gaa* KO models [24–26], all demonstrate null or nearly-zero GAA enzyme activity. Nevertheless, no neonatal mortality has been observed in any model, while neonatal death is the inevitable clinical outcome in untreated IOPD patients[27, 28]. Only one *Gaa* KO model on a DBA/2J background (homozygous *Ltbp4^Δ36^* alleles) is reported to have a shorter lifespan (but still not neonatal lethality) in male mice, compared to male *Gaa* KO mice on the C57BL/6;129 background[26]. The DBA/2J genetic background may exacerbate the severity of respiratory muscle weakness caused by *Gaa* KO deletion, leading to earlier death than is observed in other KO models[26].

Genome editing represents a new approach to the treatment of PD, compared to traditional treatments like ERT or gene therapy. A mouse that both recapitulates clinical features of human disease and harbors orthologous pathologic gene variants serves as a valuable system for the development of innovative therapies and, most importantly, studies enabling eventual clinical trials in humans. As this model undergoes in-depth validation and studies of its clinical and immune response to standard intravenous rhGAA enzyme infusions, subsequent avenues for exploration include variant rhGAA enzyme infusions, gene therapy, and CRISPR-based genomic editing. The latter approach can be performed using CRISPR “prime editing”, which is capable of targeting more than 90% of known pathogenic mutations, including the c.1935C>A transversion [29]. In addition, multiple tissues can be obtained or derived from our *Gaa^Em1935C>A^* KI mouse to investigate the potential tissue-specific efficacy of genome correction-based therapeutics *in vitro*, before *in vivo* studies are attempted. With these advances, and high sequence conservation surrounding the mutation, the *Gaa^Em1935C>A^* KI mouse represents an ideal candidate for the development of personalized therapeutics like prime editing to correct pathogenic variants, restore GAA enzyme activity and further improve functional phenotypes before translational application to the clinic.

## Materials and Methods

### *Gaa^c.1935^* guide RNA spCas9 expression vector cloning

All oligonucleotides applied in this project were manufactured by Integrated DNA Technologies (Coralville, IA). Guide RNA (gRNA) oligonucleotides with *Bbs*I (New England Biolabs) restriction enzyme overhangs were designed with forward oligo (5’-CACCG(gRNA)-3’) and reverse oligo (5’-AAAC(reverse complement gRNA)C-3’). Complementary gRNA oligonucleotides were cloned into pSpCas9(BB)-2A-Puro plasmid (pX459; Addgene plasmid ID# 48139) using the *Bbs*I I site. Positive pX459-gRNA clones were confirmed by Sanger sequencing and further expanded using the PureLink HiPure Plasmid Midiprep Kit (Invitrogen). All donor single-stranded oligodeoxynucleotides (ssODNs) were designed with 50-bp homology arms flanking the target locus and silent mutations in the <u>p</u>rotospacer <u>a</u>djacent <u>m</u>otif (PAM) as well as seed region (5 nt upstream of PAM) to prevent further Cas9 activity after successful HDR.

### *In vitro* testing of *Gaa^c.1935^* guide RNAs

pX459-*Gaa^c.1935^* gRNA expression vectors and donor ssODNs were transfected into murine C2C12 myoblast cells (ATCC CRL-1772) using the Neon™ Transfection System (ThermoFisher Scientific) as previously described[15]. In short, 3×10^5^ cells were mixed with 4.5 μg gRNA expression vector(s) and 450 nM ssODN (Table 1), then and electroporated using the following parameters: pulse voltage, 1650V; pulse width: 10 ms; pulse number: 3. Forty-eight hours posttransfection, cellular genomic DNA was obtained for Sanger sequencing around the *Gaa^c.1935^* target locus. spCas9 nuclease activity and HDR KI efficiency were determined by <u>T</u>racking of <u>I</u>ndels by <u>D</u>ecomposition (TIDE)[30] or <u>T</u>racking of <u>I</u>nsertion, <u>De</u>letions, and <u>R</u>ecombination events (TIDER)[31] analysis of DNA sequence electropherogram files.

### Generation of a *Gaa^c.1935C>A^* KI C2C12 cell line

Similar parameters to those described above were applied to transfect 4.5 μg of gRNA-1, gRNA-2 under U6 promoter in pX459 expression vectors, and 450 nM ssODN in 3×10^5^ C2C12 myoblast cells. pCMV6-AC-GFP (OriGene) was used as a positive control for transfection; and neomycin resistance gene served as negative control for puromycin selection. Electroporated cells were selected by adding 2.5 μg/mL puromycin dihydrochloride (Sigma-Aldrich) to the culture medium after 24 h of transfection. Puromycin was supplied every 48 h until all pCMV6-AC-GFP-transfected cells were no longer viable. After puromycin-resistant screening, single cell clones were selected by standard serial dilution methods in 96-well plates in the presence of 2.5 μg/mL puromycin dihydrochloride. Sanger sequencing was used to confirm the genotype of each single cell clone.

### Generation of *Gaa^Em1935C>A^* KI mice

The generation of *Gaa^Em1935C>A^* KI mice was performed at the University of California, Irvine, and all study procedures were reviewed and approved under IACUC protocol #AUP16–63. Standard methods were applied to produce pronuclear stage C57BL/6NJ embryos [16]. In brief, 3 μM crRNA/tracrRNA/3xNLS-Cas9 protein and 10 ng/μL ssODN were injected into pronuclear stage C57BL/6NJ embryos (Table 2). Surviving embryos were implanted into oviducts of 0.5dpc ICR pseudo-pregnant females.

### Whole-genome sequencing and analysis

Whole genome sequencing (WGS) and analyses were performed on tail samples from G_0_ wild type (*Gaa^wt^*), G_0_ founder #1 (*Gaac.^1935Founder#1^*), and G_0_ founder #2 (*Gaac.^1935Founder#2^*) mice. In brief, WGS was performed and analyzed on an Illumina HiSeq X Ten Sequencer at 40–50x read depth (Fulgent Genetics) using TrueSeq DNA libraries created from 1 μg fragmented genomic DNA. WGS on-target and off-target analyses were performed on the OnRamp BioInformatics platform. Data were aligned to the Mouse genome (mm10) using BWA[32]. PCR artifacts were identified with the memtest utility from Sentieon [33], and filtered out using samtools[34]. Alignments were de-duplicated and realigned around insertions and deletions using LocusCollector, Dedup, and Realigner from Sentieon. Single nucleotide variation (SNV) calling was performed with GVCFtyper from Sentieon, using the mouse dbSNP 142 data (http://hgdownload.cse.ucsc.edu/goldenpath/mm10/database/snp142.txt.gz) as the known SNPs. Known SNPs and variants falling in un-located chromosomes were removed from analysis.

For off-target analysis, we used SNVs that had C>A transversion and met the following criteria indicative of an ectopic HDR event: flanked by an N→A mutation 3 bases upstream, an N→A mutation 6 bases upstream, an N→C mutation 12 bases upstream, an N → T mutation 15 bases upstream, or an N→C mutation 39 bases upstream. This search step was repeated for the reverse complement sequences. The fully processed BAM files (after Realigner) were used as input to the Manta structural variant caller[35]. For each of the non-wild-type (WT) samples, Manta somatic caller was applied with the C57BL6-WT sample as “normal” and the sample of interest as “tumor”, thereby subtracting the background structural variants in C57BL6-WT compared to mm10. Vcf (https://vcftools.github.io) was used to annotate the output VCF files from Manta.

### Experimental Animals

The use and care of animals used in this study adhered to the guidelines of the NIH Guide for the Care and Use of Laboratory Animals, as utilized by the CHOC Children’s Institutional Animal Care and Use Committee. All study procedures were reviewed and approved under CHOC IACUC protocol #160902.

Animals were maintained and backcrossed onto a C57BL/6NJ background. Genotyping was performed by Sanger sequencing to confirm the *Gaa^c1935^* target locus with the following primers: *GAA*_c1935(F), 5’-CAGGCGTTAGGACAAATGGA-3’; *GAA*_c1935(R), 5’-TTCCAGCAGGTATGGGATTAAC-3’. Heterozygous (*Gaa^wt/Em1935C>A^*) males and females were crossed to obtain homozygous KI, heterozygous (HET), and WT mice for this study. Experiments were performed on age-matched mice of either gender (usually littermates). Homozygous knockout (KO) (B6;129-*Gaa^tm1Rabn^*/J) [24]mouse tissues for comparative molecular and biochemical analyses were acquired from Jackson Laboratory (Bar Harbor, ME).

### Quantitative real-time PCR

Total RNA was extracted from tail tip or liver homogenate using a Direct-zol RNA miniprep kit (Zymo Research) and reverse-transcribed using an iScript™ cDNA Synthesis Kit (Bio-Rad). As per the manufacturer’s instructions, both oligo(dT) and random hexamer primers were used to synthesize cDNA. The resulting cDNA was diluted 10-fold, and a 2-μl aliquot was used in a 12-μl PCR reaction with SsoAdvanced Universal Probes Supermix (Bio-Rad) and specific TaqMan primer/probe assays for *Gaa* (Taqman assay #Mm00484581_m1) and *Gapdh* (TaqMan assay #Mm99999915_g1). PCR reactions were run in triplicate and quantified with Bio-Rad CFX96 Touch Real-Time PCR Detection. *Gapdh* was used as an internal reference gene, and relative quantification of *Gaa* gene expression was measured using the comparative ΔCt method.

### Biochemical analyses

For the GAA activity assay, phosphate-buffered saline (PBS)-flushed mouse tissues or C2C12 myobloast cell pellets were homogenized in CelLytic M cell lysis reagent (MilliporeSigma). Acidic α-glucosidase enzyme activity was assessed as previously described with minor modifications[15, 36]. In brief, 10 μL tissue homogenate was mixed with 10 μL of 6 mM 4-methylumbelliferyl-α-D-glucopyranoside substrate (MilliporeSigma) in McIlvaine citrate/phosphate buffer (pH 4.3) and quenched with 180 μL glycine carbonate buffer (pH 10.5) after 1-hr incubation at 37 °C in a 96-well plate. GAA activity reactions were run in triplicate, and fluorescence measurements were obtained using an Infinite M Plex spectrofluorophotometer (Tecan) at excitation and emission wavelengths of 360 nm and 450 nm, respectively. One GAA enzymatic activity unit was defined as 1 nmol converted substrate per hour. Protein concentration was estimated using a Pierce BCA assay kit (ThermoFisher), using bovine serum albumin as a standard. Specific activity was calculated as units of GAA enzymatic activity per mg of protein.

Tissue glycogen levels were measured using a glycogen assay kit (Sigma-Aldrich) according to the manufacturer’s instructions. In brief, 10 μL tissue homogenate was incubated with hydrolysis enzyme reaction mixture in a final volume of 50 μL at room temperature for 30 min before adding 50 μL development enzyme reaction mixture for 30 min incubation at room temperature. Absorbance at 570 nm was measured using an Infinite M Plex spectrofluorophotometer (Tecan). A standard curve was generated using standard glycogen solution provided in the assay kit. Glycogen quantification assays were performed in duplicate, and an extra reaction without hydrolytic enzyme treatment was used for background correction of endogenous glucose levels in each sample. Tissue glycogen level is expressed as μg of glycogen per mg of protein.

### LC3B western blot analysis

Frozen mouse tissues were homogenized in CelLytic M cell lysis reagent (MilliporeSigma) and cOmplete protease inhibitors (Roche) was added to prevent protein degradation. Total protein concentration of the supernatants from centrifuged tissue lysates was determined by BCA protein assay (Pierce). Eight micrograms of total protein lysate were resolved on 4–15% Mini-PROTEAN TGX Stain-free gels (Bio-Rad) and transferred onto Immuno-Blot PVDF membranes (Bio-Rad). Membrane blots were blocked with EveryBlot blocking buffer (Bio-Rad) and probed with an anti-LC3B primary antibody (rabbit polyclonal, Sigma L7543, 1:1000 dilution) followed by an HRP-conjugated secondary antibody (Bio-Rad 1706515; 1:3000 dilution) before applying ECL HRP substrate (Bio-Rad 1705060) for chemiluminescence. Stain-free gels and blots were imaged using the stain-free and chemiluminescence settings on the ChemiDoc™ MP imaging system (Bio-Rad). LC3B-I and LC3B-II protein levels were measured by densitometric analysis of western blots using Fuji software (ImageJ version 2.0)[37]. Signals were normalized to the amount of total protein as determined by densitometric analysis of stain-free gels. Absolute ratios of LC3B-II to total LC3B were calculated as follows: LC3B-II/(LC3B-I + LC3B-II)).

### Murine echocardiography

Transthoracic echocardiography (M-mode and 2-dimensional echocardiography) was performed using a Vevo 2100 high-resolution ultrasound system, with a linear transducer of 32–55 MHz (VisualSonics Inc.). Chest fur was removed by using depilatory cream one day prior to the procedure. Mice were kept warm on a heated platform (37 °C) and anesthetized with 5% isoflurane for 15 seconds, then maintained at 0.5% throughout the echocardiography examination. Small needle electrodes for simultaneous electrocardiography were inserted into one upper and one lower limb. Measurements of chamber dimensions and wall thickness were performed while heartbeats of the mice were greater than 500 beats per minute (bpm). Percentage fractional shortening (%FS) was used as an indicator of left ventricular systolic cardiac function and calculated as follows: %FS = (LVIDd – LVIDs)/LVIDd * 100.

### Forelimb grip strength assay

Forelimb grip strength was measured as previously described[38]. Following acclimatization (at least one hour prior to grip strength measurement), each mouse was weighed and placed on a forelimb pull bar attached to an isometric force transducer (Columbus Instruments, Columbus, OH, USA). The mouse was pulled away from the bar by its tail, and the force required was recorded by the force transducer. Over 3 consecutive days, each mouse performed 3 pulls per day for a total of 9 pulls per test session. Peak tension force (N) was calculated as the average of each subject’s 9 pulls over the test session and normalized by body weight.

### Tissue harvesting, processing, and histological staining

Three-month-old mice were euthanized using CO_2_ asphyxiation and transcardially perfused with PBS. Brains were dissected sagittally along the midline; left hemispheres were rapidly frozen and stored at −80 °C for biochemical analysis, and right hemispheres were post-fixed at 4°C in zinc formalin. Heart, diaphragm, and gastrocnemius muscle were also harvested. Half of the tissue samples for biochemical studies were rapidly frozen and the other half of tissues were post-fixed at 4°C in zinc formalin.

Samples for histological staining were processed and embedded in paraffin blocks for sectioning at 4-μm thickness, and periodic acid-Schiff (PAS) staining (Sigma-Aldrich) was performed according to the manufacturer’s instructions. EVOS M5000 imaging system (Invitrogen) was used to capture representative images at 20x objective magnification on RGB-mode illumination. To quantify PAS staining intensity, the RGB channel of PAS staining was used to determine the correct optical density (OD) vectors according to the protocol previously described[39, 40]. In brief, images with homogeneous background from three muscle tissues (heart, diaphragm, and gastrocnemius), and three different areas of brain (frontal cortex, hippocampus, and cerebellum), were used for analysis. The background-corrected images were prepared with commonly used color deconvolution with defined color vectors[40] using Fuji software (ImageJ version 2.0)[37]. Images were then color-convoluted using the H-PAS function. Image 2 (PAS) was converted to 8-bit, and data were presented as OD (mean grey value) as calculated with the following formula: OD = log (max intensity/mean intensity), where max intensity is 255 for 8-bit images. Average mean OD was compared between groups.

### Statistical analysis

All graphs and statistical comparisons were generated using GraphPad Prism 9. Statistical analyses were performed using the two-tailed unpaired *t*-test or one-way ANOVA followed by Tukey’s HSD test. All data are presented in this study as mean ± standard deviation (SD).

## Acknowledgements

This work was supported by the UCLA Intercampus Medical Genetics Training Program T32 (T32GM008243), the Chao Family Comprehensive Cancer Center Transgenic Mouse Facility Shared Resource, supported by the National Cancer Institute of the National Institutes of Health under award number P30CA062203, the Campbell Foundation of Caring, The Larry and Helen Hoag Foundation, and a CHOC OneWish Grant. The content is solely the responsibility of the authors and does not necessarily represent the official views of the National Institutes of Health.

## Author Contribution

Conceived and designed the experiments: SK, JYH, RYW

Performed the experiments: SK, JYH, JH, AR, NDD, CC, YC, JN, RYW

Analyzed the data: SK JYH, JH, AR, NDD, YC, JD-T, RYW

---

**Supplementary Figure 1: No gender difference in anatomical features of left ventricular cardiac hypertrophy in 3-month-old *Gaa^c.1935C>A^* knock-in mice.** Comparison of ventricular measurements and myocardial contraction in (A) male and (B) female mice. All measurements from WT (n=12; 6F, 6M), HET (n=10; 5F, 5M), and KI (*Gaa^.Em1935C>A^*; n=10; 5F, 5M) mice. Data are shown as mean ± SD. All comparisons were analyzed using one-way ANOVA with the Tukey post-hoc test. \**p*<0.05, \*\**p*<0.01, \*\*\**p*<0.001, \*\*\*\**p*<0.0001, ns: not significant.

**Supplementary Figure 2: PAS and PAS-D staining showing glycogen storage in *Gaa^c.1935C>A^* knock-in mouse tissues.** Top rows are the same images shown in Figure 7 in the main text. Bottom rows are adjacent slide sections of the same tissue showing PAS-D staining in (A) muscle tissues and (B) brain sections. (A) Abnormal glycogen accumulation in cardiac and skeletal muscle tissues was observed in KI mice compared to WT mice (top). Staining is no longer seen after diastase treatment, providing evidence for glycogen accumulation. Arrowheads indicate regions of glycogen accumulation. (B) No significant difference in PAS or PAS-D staining in brain sections was observed between WT and KI mice. Scale bar represents 100 μm.

