## Supplementary figures and images for "CRISPR-Mediated Generation and Characterization of a *Gaa* Homozygous c.1935C>A (p.D645E) Pompe Disease Knock-in Mouse Model Recapitulates Human Infantile Onset-Pompe Disease"

### Supplementary Figure 1

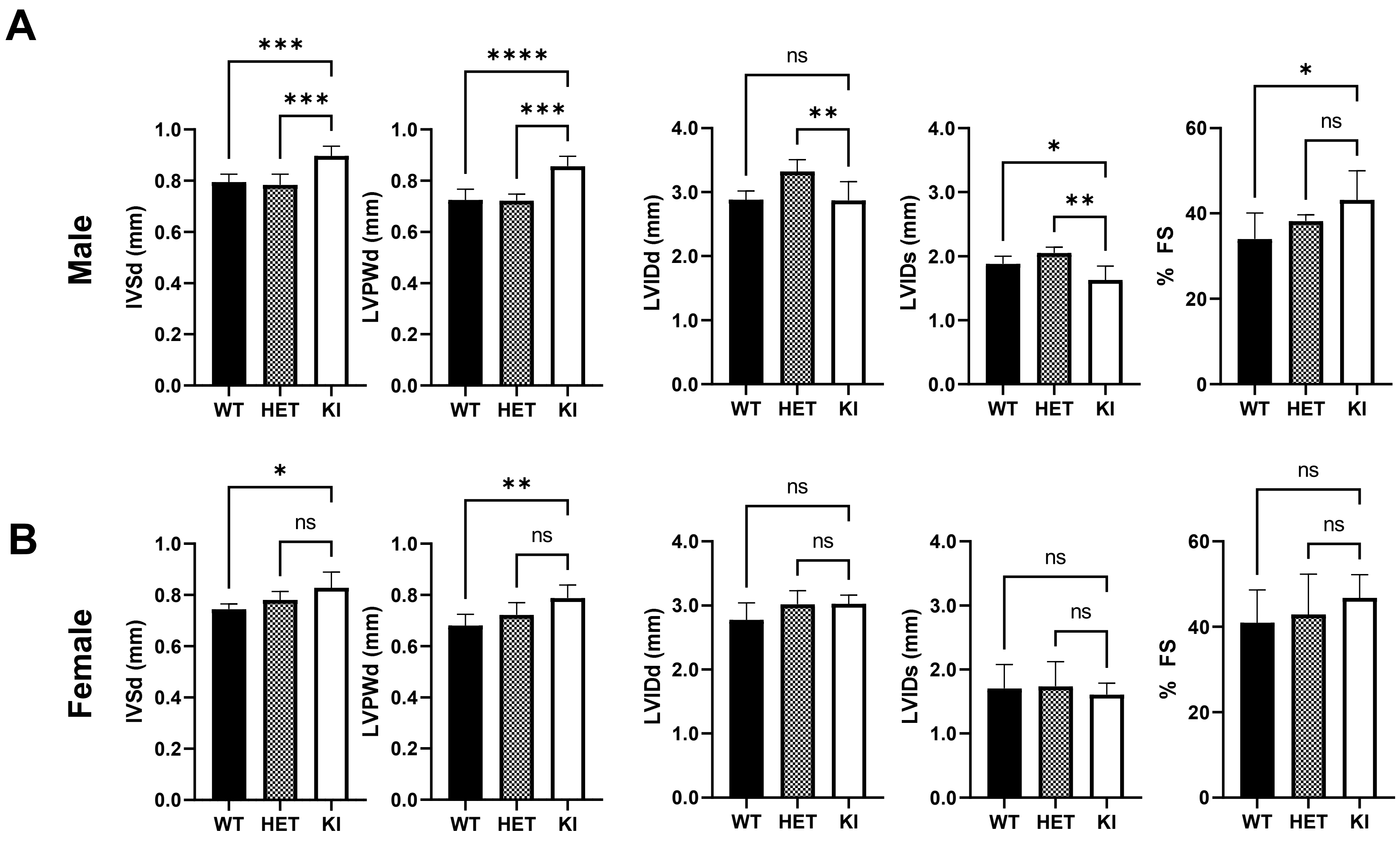

### Supplementary Figure 2

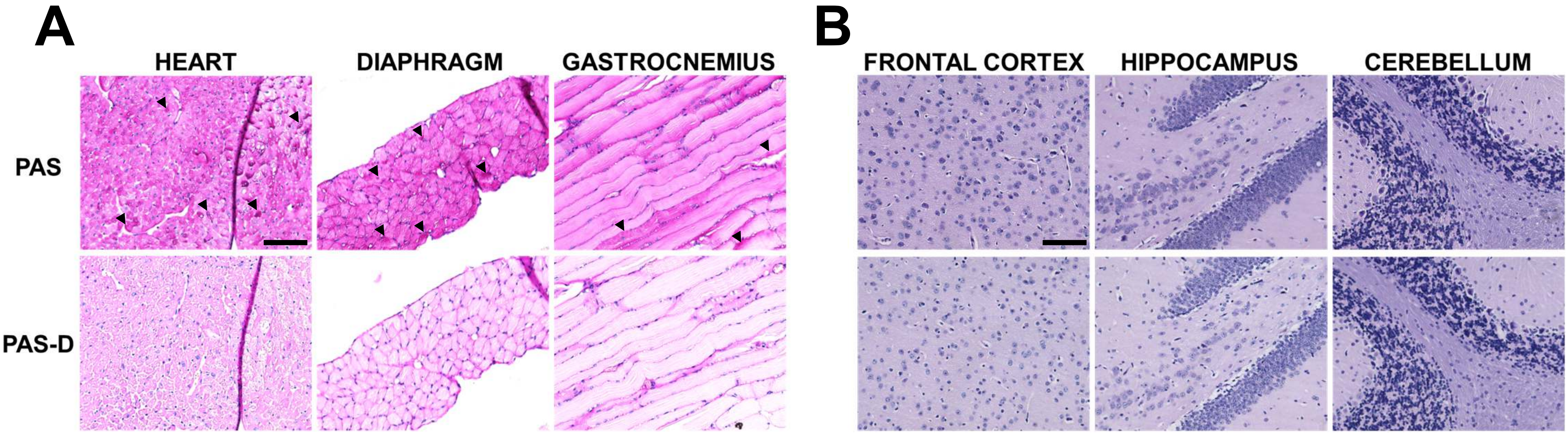
